# Genes for 5’mRNA cap binding proteins, *eif4ea* and *eif4eb*, have alternative roles during heart regeneration

**DOI:** 10.1101/2025.05.27.656424

**Authors:** Rejenae Dockery, Mateo Zevallos, Carson McNulty, Tessa Zecchino, Reece Ratermann, Richard White, Joseph Aaron Goldman

## Abstract

All eukaryotes possess the essential mRNA cap-binding protein eIF4E, which initiates translation. This protein is highly conserved, with versions from zebrafish and humans able to rescue yeast. All fish also have an additional eIF4E family, called *eif4e1c*, which is crucial for heart regeneration. Deletion of *eif4e1c* leads to growth defects and impaired heart regeneration, but it’s unclear if similar effects occur when canonical translation mechanisms are disrupted. In this study, we deleted the two zebrafish paralogs of eIF4E, *eif4ea* and *eif4eb*. While individual deletions of these genes do not show the same phenotypes as *eif4e1c* deletion, compound mutants enhanced *eif4e1c* phenotypes, suggesting partial compensation by canonical eIF4E proteins in *eif4e1c* mutants. Surprisingly, unlike other eukaryotes, deleting both canonical mRNA cap-binding proteins did not result in lethality, and *eif4e1c* could fully compensate for eIF4E function. Although we anticipated that double mutants would regenerate hearts better since they contain *eif4e1c* alone, no improvement was observed in the double mutant. Interestingly, single deletions of *eif4ea* or *eif4eb* did improve heart regeneration. This supports a model where the balance between the *eif4e1c* and canonical pathways is crucial for stimulating heart regeneration.

## INTRODUCTION

The rate determining step for the initiation of translation is binding of eIF4E to the 5’ cap of the mRNA^1,2^. Most of what is known about eIF4E comes from *in vitro* studies of purified components or in many cases from cell culture. *In vivo* studies are challenging since eIF4E orthologs are required for viability from yeast to mouse^3,4^. Only a few studies have been able to determine loss-of-function phenotypes for eIF4E in an organism. In C. *elegans*, knockout of *ife*-2 results in an extended lifespan but with few other observable phenotypes^5^. The presence of a second ortholog (*ife*-4) that is expressed organism-wide is likely compensating unlike in yeast and mice where only one canonical cap-binding gene is present. In addition to eIF4E, several paralogs have emerged from vertebrate-specific whole genome duplications^6^. eIF4E2 and eIF4E3, which have only 26-35% sequence identity with eIF4E, function in specific tissues or contexts and often inhibit rather than initiate translation^6–9^. Another paralog, eIF4E1B, is 66-75% identical to eIF4E and is exclusively present in germ cells, and mouse knockouts have shown that it protects maternal mRNA from degradation before the zygotic genome is activated^10,11^. How different mRNAs are selected by each cap binding protein and what determines their function is an area of active research^12^. To date, none of the eIF4E orthologs have been shown to be capable of performing canonical functions.

Zebrafish possess all the different types of cap-binding proteins that have been described, including two genes for the canonical eIF4E protein, named *eif4ea* and *eif4eb*. Additionally, zebrafish have a unique family of cap-binding homologs, called Eif4e1c, which is found only in fish and not in any terrestrial vertebrates^13^. The highly conserved amino acids necessary for binding to the cap and initiating assembly of the ribosome complex are also present in the Eif4e1c family^14,15^. Moreover, there are 23 amino acids (13% of the total protein) that are shared among all Eif4e1c proteins with remarkable evolutionary conservation. For example, the Eif4e1c proteins in sharks and zebrafish are 86% identical and 95% similar, with only 8 of 186 significant amino acid differences over ∼480 million years of evolution. Thus, while the Eif4e1c family shares features with the canonical eIF4E in critical cap binding residues, Eif4e1c is likely a unique homolog with highly conserved and distinctive properties.

Previously, we showed that knockout of *eif4e1c* leads to decreased survivability, less growth and impaired cardiac regeneration. Such profound negative consequences had not previously been described for mutants in 5’mRNA cap binding proteins. While deletion is lethal, the dosage of eIF4E can be reduced significantly with little obvious effects^5,16^. An acute 90% reduction of eIF4E protein in yeast led to an initial 75% reduction in translation that was largely recovered after 4 hours^17^. It remains unclear if *eif4e1c* is functioning as an interchangeable third paralog to *eif4ea* and *eif4eb* or if *eif4e1c* is dedicated to specific pathways. Expression levels of canonical eIF4E can impact target mRNA binding. For example, *Eif4e* mutant mice that are heterozygous do not show significant phenotypes, except for an increased resistance to cancer^18^. When *Eif4e* is present at half the normal dose, a subset of mRNAs related to oxidative stress are translated less efficiently. Thus, it remains possible that phenotypes may arise in Δ*eif4e1c* mutants when total cap-binding protein levels or activity drops below a required threshold. Further experiments are necessary to clarify.

To see if growth and survival phenotypes in Δ*eif4e1c* mutants are a general feature of inhibiting translation, we established stable mutant alleles of *eif4ea* and *eif4eb* in zebrafish using CRISPR. The knockout of each individual canonical gene results in homozygous mutant fish at Mendelian ratios and we didn’t detect any size differences in either of the single mutants. Surprisingly, even though canonical factors are generally essential, a double knockout of *eif4ea* and *eif4eb* has normal viability and growth. This suggests both that Eif4e1c is perfectly capable of compensating for canonical translation and that Δ*eif4e1c* phenotypes are pathway specific. However, phenotypes did emerge in Δ*eif4ea* and Δ*eif4eb* single mutants during regeneration. While heart regeneration in Δ*eif4ea* or Δ*eif4eb* mutants improves, regeneration of the fins is worse.

## RESULTS

### The *eif4e1c* gene can fully compensate for loss of *eif4ea* and *eif4eb*

To determine whether mRNA cap binding proteins are generally required for survival and proper growth, we used CRISPR-mediated mutagenesis to create deletion mutants for both zebrafish canonical paralogs, *eif4ea* and *eif4eb*. Using two sgRNA we removed 4,262bp from the *eif4ea* (Δ*eif4ea*) gene, deleting 65% of the central part of the protein (Figure 1A, highlighted red). For *eif4eb* (Δ*eif4eb*), the two sgRNA removed 560bp including 21% of the coding sequence and the required cap-binding residue, Trp-54 (Figure 1B, highlighted red). Both mutations are expected to be null mutations. When heterozygotes were crossed, the individual mutations were found at Mendelian ratios for both Δ*eif4ea* and Δ*eif4eb* (Table 1). Therefore, unlike with *eif4e1c*, mutants for *eif4ea* and *eif4eb* have normal viability.

**Figure 1.**
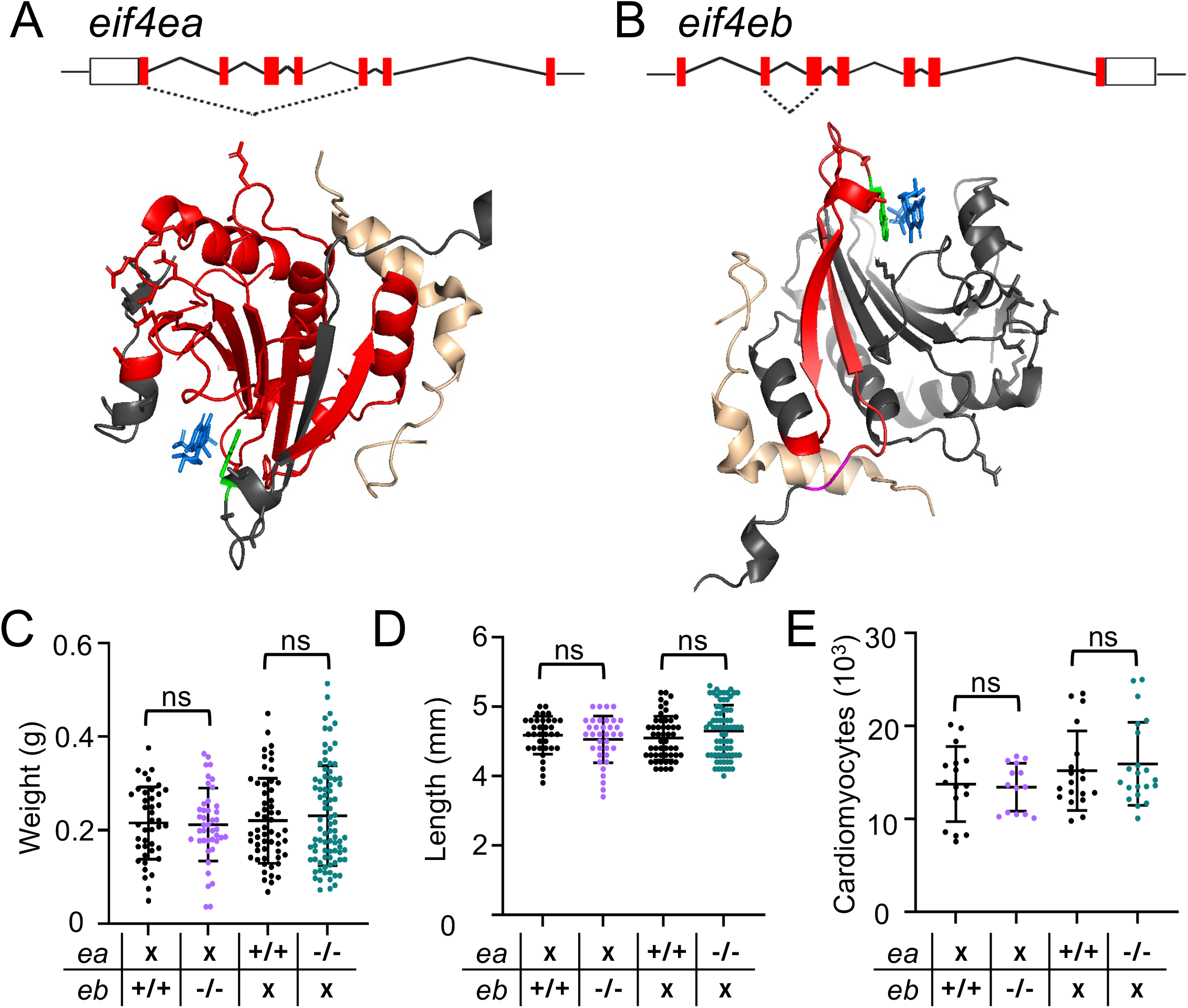
Deletion of genes for canonical cap-binding proteins results in no impairment to zebrafish survival or growth. **(A)** A 4,262bp region from exon 2 to exon 7 was removed from the *eif4ea* locus using CRISPR. The region deleted from the protein is highlighted in red in the crystal structure. **(B)** A 562bp region from exon 2 to exon 3 was removed from the *eif4eb* locus suign CRISPR. The region deleted from the protein (red) includes the residues required for cap binding (green). In both crystal structures a cap analog is present and highlighted in blue. **(C)** Weights of wildtype (black, first column) and Δ*eif4ea* siblings (purple) are shown to be similar (wild type mean = 0.215g, Δ*eif4ea* mutant mean = 0.212g, Welch’s t-test p-value = 0.843, N = 41,42). Weights between Δ*eif4eb* (green) and their wildtype siblings (black, third column) were also found to be the same (wild type mean= 0.220g, Δ*eif4eb* mean = 0.231g, p value = 0.538, N=56,84). **(D)** Length of Δ*eif4ea* (purple) and Δ*eif4eb* (green) are also the same as wildtype siblings (Mean: wild type = 25.89mm, Δ*eif4ea* = 25.28mm, wild type = 25.47mm, Δ*eif4eb* = 26.48mm; p value = 0.407,0.117; N = 35, 36, 51, 73). **(E)** The two mutant fish also had no changes in the numbers of CMs (Mean: wild type = 2748, Δ*eif4ea* = 2682, wild type = 3038, Δ*eif4eb* = 3184; p value = 0.850, 0.601; N = 16, 14, 19, 21).

To determine whether these mutations lead to growth deficits, we intercrossed heterozygotes for Δ*eif4ea* and grew them to adulthood. The homozygous Δ*eif4ea* adult mutants weighed the same as their wild type siblings (Figure 1C, wild type mean = 0.215g, Δ*eif4ea* mutant mean = 0.212g, Welch’s t-test p-value = 0.843, N = 41,42). The homozygous Δ*eif4eb* adult mutants also weighed similarly to wild type (wild type mean= 0.220g, Δ*eif4eb* mean = 0.231g, Welch’s t-test p value = 0.538, N=56,84). The lengths of both mutants were nearly identical to wild type (Figure 1D; wild type mean = 25.89mm, Δ*eif4ea* mean = 25.28mm, wild type mean = 25.47mm, Δ*eif4eb* mean = 26.48mm; Welch’s t-test p value = 0.407 and 0.117; N = 35, 36, 51, 73). We conclude that unlike mutants in their ortholog *eif4e1c,* individual mutants of *eif4ea* and *eif4eb* do not demonstrate growth deficits.

Previously, we showed that the *eif4e1c* mutants had overall weight deficits that were reflected by reduced cell number^13^. A similar deficit might be occurring in the canonical mutants that may be masked when only measuring the overall weight. We removed adult hearts and counted the numbers of cardiomyocytes (CMs) as was done for *eif4e1c*. However, similar numbers of CMs were found in both canonical mutants when compared to their wild type siblings (Figure 1E; wild type mean = 2748, Δ*eif4ea* mean = 2682, wild type mean = 3038, Δ*eif4eb* mean = 3184; Welch’s t-test p value = 0.850, 0.601; N = 16, 14, 19, 21). Thus, canonical mutants have similar cell numbers, at least in the heart, suggesting that they do not recapitulate any of the phenotypes observed in *eif4e1c* mutants.

The two zebrafish paralogs of canonical eIF4E are 86% identical (95% in the core), and they likely readily compensate for one another. Total loss of canonical eIF4E is lethal in eukaryotes, so we did not expect to recover Δ*eif4ea /* Δ*eif4eb* double mutants. Yet, we set up a Δ*eif4ea /* Δ*eif4eb* heterozygotes intercross to evaluate possible mutant and heterozygote combinations for Δ*eif4e1c*-like phenotypes. To our surprise, adult double homozygous Δ*eif4ea /* Δ*eif4eb* mutants were recovered at Mendelian ratios (Figure 2A, χ^2^ = 8.55, p = 0.38). Furthermore, they did not demonstrate any significant difference in weight or length and hearts had similar numbers of CMs (Figure 2B-D, weight – ANOVA p-value = 0.085, length – ANOVA p-value = 0.130, CMs Welch’s t-test p-value = 0.11). Unlike in other eukaryotes, complete deletion of canonical mRNA cap binding proteins in zebrafish results in viable adults without any observable phenotypes. These results favor a model where *eif4e1c* is a specialized translation initiation factor and its unique function is what leads to survival and growth phenotypes in Δ*eif4e1c*.

**Figure 2.**
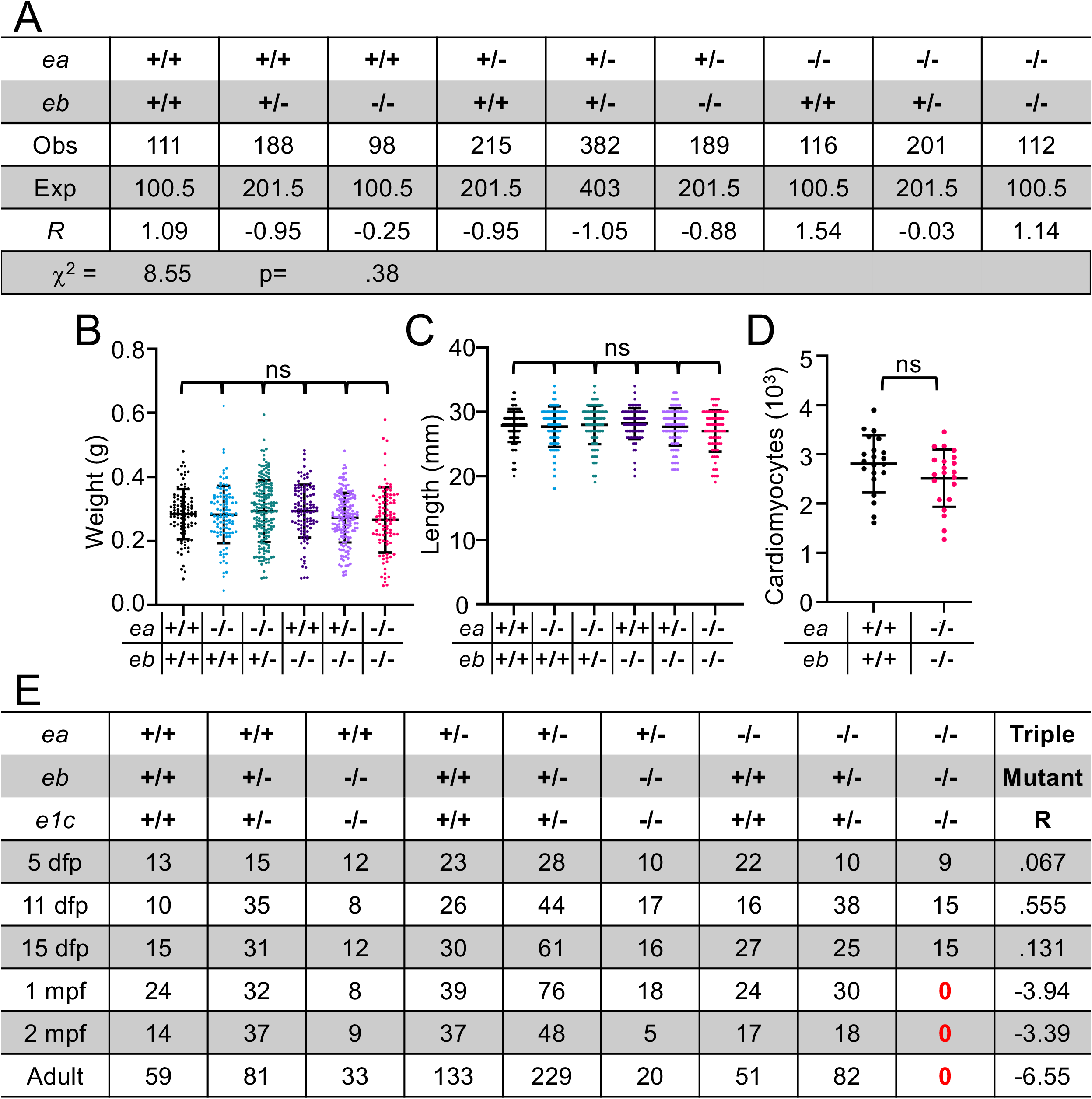
The *eif4e1c* gene can fully compensate for complete loss of both canonical cap binding orthologs. **(A)** Table shows survival of Δ*eif4ea /* Δ*eif4eb* double mutants at Mendelian ratios. *R* is the critical value and the Δ^2^ statistics are at the bottom. **(B)** Weights of adult siblings that are Δ*eif4ea* or Δ*eif4eb*. There is no significant difference (ANOVA p-value = 0.085). **(C)** Lengths of adult siblings that are Δ*eif4ea* or Δ*eif4eb* compared to wildtype. There is no significant difference (ANOVA p-value = 0.130). **(D)** CM numbers of wildtype (black) and Δ*eif4ea /* Δ*eif4eb* double mutants (pink) are not significant (Welch’s t-test p-value = 0.11). **(E)** Time course of triple mutant survival. The Δ*eif4ea /* Δ*eif4eb* double mutants survive at Mendelian ratios into adulthood yet the **triple mutants** die 15-30 days post fertilization. Thus, Δ*eif4ea /* Δ*eif4eb* survival depends on *eif4e1c*.

To determine whether Δ*eif4ea /* Δ*eif4eb* double mutants survive on *eif4e1c* alone we intercrossed Δ*eif4ea /* Δ*eif4eb /* Δ*eif4e1c* triple heterozygotes to look for triple mutants. As expected, no triple homozygotes were recovered as adults (Table 1). Previously we showed that the Δ*eif4e1c* solo mutants die between 4- and 8-weeks post fertilization (wpf) at juvenile stages^13^. To determine when the triple homozygotes die, we performed a time course. Triple homozygotes were found at expected levels at 5-, 11-, and 15-days post fertilization (dpf) (Figure 2E, standardized residual = 0.067, 0.555, 0.131). However, by 4wpf, no triple mutants were recovered (Figure 2E, standardized residual = -3.94). Thus, without compensation by canonical alleles, the Δ*eif4e1c* mutants die sooner. Likely, the Δ*eif4e1c* embryos survive until 15-30dpf due to maternal deposition of mRNA and protein for all three genes. This data also shows that Δ*eif4ea /* Δ*eif4eb* double mutants are fully compensated by the presence of *eif4e1c* alone. Thus, unlike other eIF4E orthologs, Eif4e1c is a cap binding protein that is likely to be fully functional and capable of canonical translation initiation.

### Both canonical *eif4ea* and *eif4eb* each partially compensate for Δ*eif4e1c* mutant phenotypes

The phenotypes observed in Δ*eif4e1c* fish appear to be incomplete, for example, while many mutants die as juveniles, others survive well into adulthood. Furthermore, increased expression of canonical protein suggests that there may be some compensation of *eif4e1c* phenotypes by canonical pathways^13^. To determine if canonical mRNA cap binding proteins partially compensate for Δ*eif4e1c* phenotypes we intercrossed *eif4ea* / *eif4e1c* double heterozygotes to make compound mutants. The combined mutation of *eif4ea* and *eif4e1c* potentiated death in Δ*eif4e1c* mutants with only 24% of the double mutants reaching adulthood (Figure 3A). The Δ*eif4e1c* single mutants; however, were not significantly underrepresented by Chi square analysis suggesting that the phenotypes may be waning with subsequent generations (95 vs 107). Interestingly, even loss of a single *eif4ea* allele reduces survival with 30% of the Δ*eif4ea-*heterozygote */* Δ*eif4e1c-*homozygotes dying (Figure 2A, standardized residual = -4.37 vs -7.73). This shows that reducing the dosage of the *eif4ea* gene has scalable effects on the survival of *eif4e1c* mutants.

**Figure 3.**
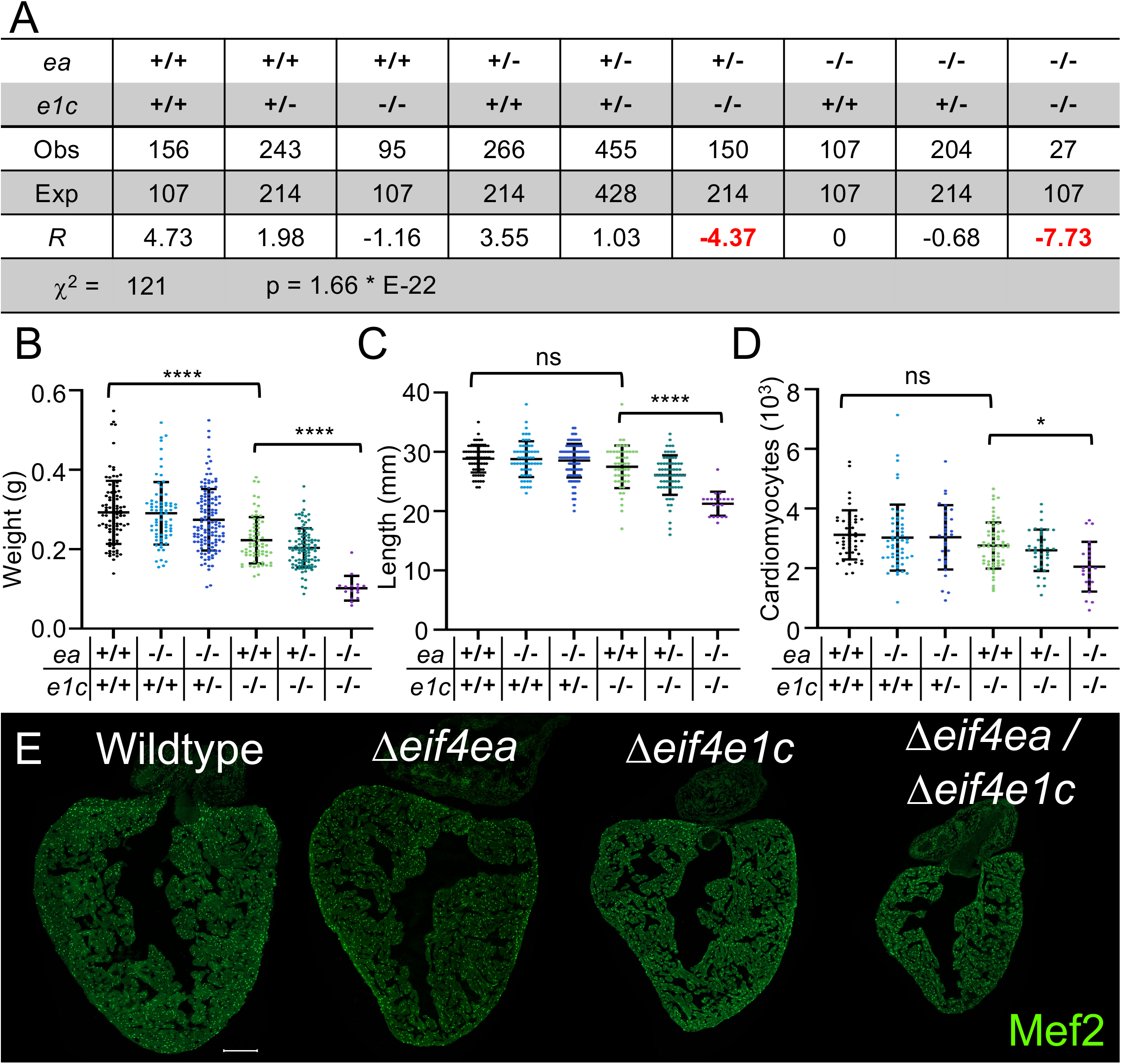
The canonical *eif4ea* partially compensates for Δ*eif4e1c* phenotypes. **(A)** Δ*eif4ea /* Δ*eif4e1c* double mutants are at 24% the level of single Δ*eif4e1c*. **(B)** The Δ*eif4ea /* Δ*eif4e1c* double mutants weigh (purple) 55% less than the Δ*eif4e1c* singles (light green) and 65% less than their wildtype (black) siblings (Mean: wildtype = 0.293g, Δ*eif4e1c* = 0.223g, Δ*eif4ea /* Δ*eif4e1c* = 0.102g, ANOVA p-value < 0.0001, N=95, 66, 16). **(C)** The double mutants are also shorter (Mean: wildtype = 28.8mm, Δ*eif4e1c* = 27.5mm, Δ*eif4ea /* Δ*eif4e1c* = 21.3mm, ANOVA p-value < 0.0001, N=95, 66, 20). **(D, E)** double mutant hearts have 66% the number of CMs versus wildtype siblings and 74% the number of CMs as Δ*eif4e1c* siblings (Mean: wildtype = 3122, Δ*eif4e1c* = 2763, Δ*eif4ea /* Δ*eif4e1c* = 2055, ANOVA p-value < 0.0001, N = 47, 61, 23).

To determine if *eif4e1c* growth deficits were similarly potentiated by mutating *eif4ea*, we weighed the adult double mutants. As expected, Δ*eif4e1c* mutants weighed 24% less than their wild type siblings (Figure 3B, wild type mean = 0.293g, Δ*eif4e1c* mean = 0.223g, ANOVA p-value < 0.0001, N=95, 66). The double Δ*eif4ea /* Δ*eif4e1c* mutants were even smaller, weighing 55% less than the Δ*eif4e1c* singles and 65% of wildtype (Figure 3B, Δ*eif4ea /* Δ*eif4e1c* mean = 0.102g, ANOVA p-value < 0.0001, N=95, 16). The lengths of the *eif4e1c* solo mutants were not significantly smaller (Figure 3C, wild type mean = 28.8mm, Δ*eif4e1c* mean = 27.5mm, ANOVA p-value = 0.150, N = 63, 49). However, previously, we did observe that this phenotype waned with age. The Δ*eif4ea /* Δ*eif4e1c* mutants were 65% shorter than wild type siblings, demonstrating that deletion of *eif4ea* further exacerbated *eif4e1c* related growth deficits (Δ*eif4ea /* Δ*eif4e1c* mean = 21.3mm, ANOVA p-value < 0.0001, N = 63, 20). The growth deficiencies are likely a result of fewer cell numbers. The Δ*eif4ea /* Δ*eif4e1c* double mutant hearts were smaller than the Δ*eif4e1c* mutants alone (Figure 3DE). There were 26% fewer CMs in the Δ*eif4ea /* Δ*eif4e1c* mutant hearts compared to Δ*eif4e1c* alone (Figure 3D, wild type mean = 3122, Δ*eif4e1c* mean = 2763, Δ*eif4ea /* Δ*eif4e1c* mean = 2055, ANOVA p-value < 0.0001, N = 47, 61, 23). Thus, we conclude that the canonical *eif4ea* gene partially compensates for Δ*eif4e1c* mutant growth deficiencies.

Since *eif4ea* compensates the phenotypes in Δ*eif4e1c* mutants, we wanted to know if this was a general feature of canonical cap-binding proteins. Therefore, we intercrossed Δ*eif4eb /* Δ*eif4e1c* heterozygotes to make compound mutants. Since the two genes are on the same chromosome, we were unable to directly compare double mutant versus individual single mutant siblings. The double mutant Δ*eif4eb /* Δ*eif4e1c* were significantly underrepresented, and their wild-type siblings were overrepresented (Figure 4A; 293 +/+: 500 -/+: 185 -/-; χ^2^=24.35, p<0.0001). However, this is not appreciably more death than we reported for Δ*eif4e1c* alone with 76% of the mutants reaching adulthood^13^. This is in stark contrast to the Δ*eif4ea /* Δ*eif4e1c* compound mutants where embryo survival is drastically attenuated to 24%. This suggests that *eif4eb* minimally compensates for the limited survival of *eif4e1c* mutants. A single copy of the *eif4ea* gene is sufficient to allow for growth to adulthood, with a slight increase in death in Δ*eif4ea* heterozygotes / Δ*eif4eb* Δ*eif4e1c* homozygotes compared to Δ*eif4eb /* Δ*eif4e1c* homozygotes alone (Figure 2E, R=-3.50, R =-1.52).

**Figure 4.**
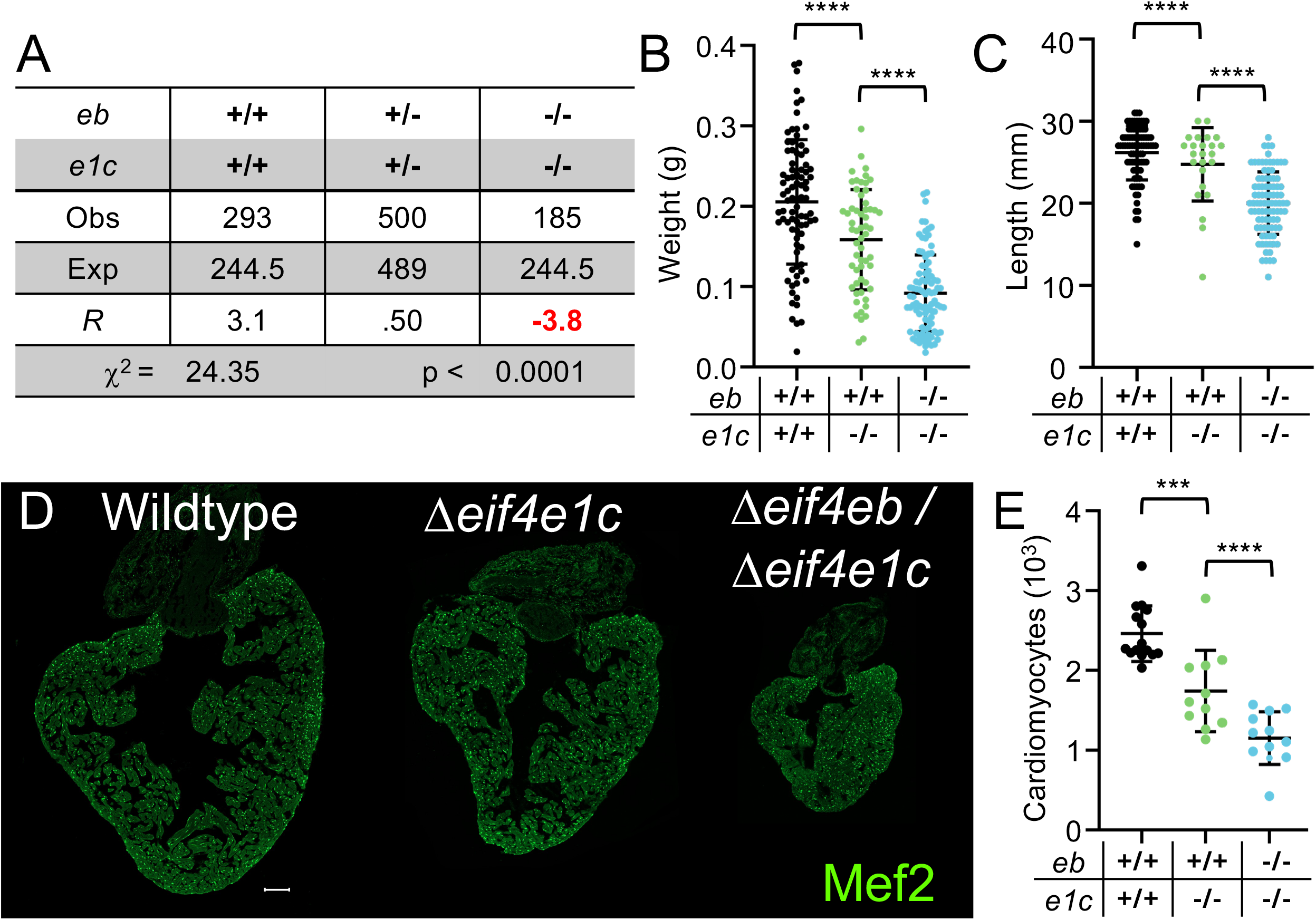
The canonical *eif4eb* compensates from growth deficits in Δ*eif4e1c*. **(A)** Adult (3-4mpf) Δ*eif4eb /* Δ*eif4e1c* double mutants are not underrepresented any more than Δ*eif4e1c* single mutants. **(B)** The Δ*eif4eb /* Δ*eif4e1c* double mutants weigh (light blue) 40% less than the Δ*eif4e1c* singles (light green) and 57% less than their wildtype (black) siblings (mean = 0.215 g, 0.159 g, 0.092 g, ANOVA p-value < 0.0001, N = 107, 58, 91). **(C)** The double mutants are also shorter (mean = 26.6mm, 24.6mm, 20.0mm, ANOVA p-value = 0.0024). **(D, E)** Δ*eif4eb /* Δ*eif4e1c* double mutant hearts have 34% the number of CMs versus wildtype siblings and 66% the number of CMs as the age-matched Δ*eif4e1c* control group (mean = 2461, 1740, 1151, ANOVA p-value < 0.0001, N = 15, 11, 12). Since *eif4eb* and *eif4e1c* are on the same chromosome individual mutants aren’t recovered.

To determine if *eif4eb* compensates for *eif4e1c* related growth deficits, we measured adult fish from the Δ*eif4eb /* Δ*eif4e1c* heterozygote cross. Since no Δ*eif4e1c* solo mutants are possible, we used an age-matched Δ*eif4e1c* heterozygote intercross as a control (see Methods). As expected, Δ*eif4e1c* mutants weighed 27% less than their wild type siblings (Figure 4B, Mean: wild type = 0.215 g, Δ*eif4e1c* = 0.159 g, ANOVA p-value < 0.0001, N = 107, 58). The double Δ*eif4eb /* Δ*eif4e1c* mutants were even smaller, weighing 40% less than the Δ*eif4e1c* singles and 57% less than their wildtype siblings (Figure 4B, Mean: Δ*eif4eb /* Δ*eif4e1c* = 0.092 g, ANOVA p-value < 0.0001, N = 91). The lengths of the *eif4e1c* solo mutants were also significantly smaller than wildtype (Figure 4C, Mean: wild type = 26.6mm, Δ*eif4e1c* = 24.6mm, ANOVA p-value = 0.0024, N = 107, 58). The double mutants were even shorter, only 75% of their wildtype siblings (Figure 4C, Mean: Δ*eif4eb /* Δ*eif4e1c* = 20.0mm, ANOVA p-value < 0.0001, N = 91). As expected, Δ*eif4eb /* Δ*eif4e1c* double mutant hearts were significantly smaller with only 34% the number of CMs versus their wildtype siblings and 66% the number of CMs as the age-matched Δ*eif4e1c* control group (Figure 4DE, Mean: wild type = 2461, Δ*eif4e1c* = 1740, Δ*eif4eb /* Δ*eif4e1c* = 1151, ANOVA p-value < 0.0001, N = 15, 11, 12). Thus, like *eif4ea*, the *eif4eb* gene compensates for Δ*eif4e1c* growth deficits.

There were also significant size differences between siblings in the Δ*eif4ea /* Δ*eif4eb /* Δ*eif4e1c* triple heterozygote intercross. Recapitulating what was shown above, there was no difference between Δ*eif4ea* mutants and their wildtype siblings but the Δ*eif4eb /* Δ*eif4e1c* double mutants were significantly smaller (Figure 5A, Mean: wildtype = 0.287g, Δ*eif4ea* = 0.315g, Δ*eif4eb /* Δ*eif4e1c* = 0.110; ANOVA wt - Δ*eif4ea* p-value = 0.712, ANOVA wt-Δ*eif4eb /* Δ*eif4e1c* p value < 0.0001; N = 47, 37, 34). Heterozygotes for Δ*eif4ea* that were also Δ*eif4eb /* Δ*eif4e1c* double mutants were even smaller, 35% the mass of even the Δ*eif4eb /* Δ*eif4e1c* double mutants (Δ*eif4ea*-heterozygotes /Δ*eif4eb /* Δ*eif4e1c* = 0.039, ANOVA p value < 0.0001, N = 14). The same differences were observed for their lengths (Figure 5B, Mean: wildtype = 27.5mm, Δ*eif4eb /* Δ*eif4e1c* = 20.5mm, Δ*eif4ea*-heterozygotes */*Δ*eif4eb /* Δ*eif4e1c* = 15.1mm, ANOVA p value < 0.001, N = 48, 34, 15). Fish with a single copy of *eif4ea* as the only cap-binding protein had 53% the number of CMs as siblings with two copies of *eif4ea* (Figure 5CD, Mean: wildtype = 2882, Δ*eif4eb /* Δ*eif4e1c* = 1962, Δ*eif4ea*-heterozygotes */*Δ*eif4eb /* Δ*eif4e1c* = 1046, ANOVA p value < 0.0001, N = 9, 10, 9). This demonstrates a clear dosage effect where the number of alleles of wildtype canonical factors proportionally impacts phenotypes associated with the loss of *eif4e1c*. Altogether, 47% of zebrafish with a single copy of *eif4ea* survive, but they are 14% the size of their wildtype siblings.

**Figure 5.**
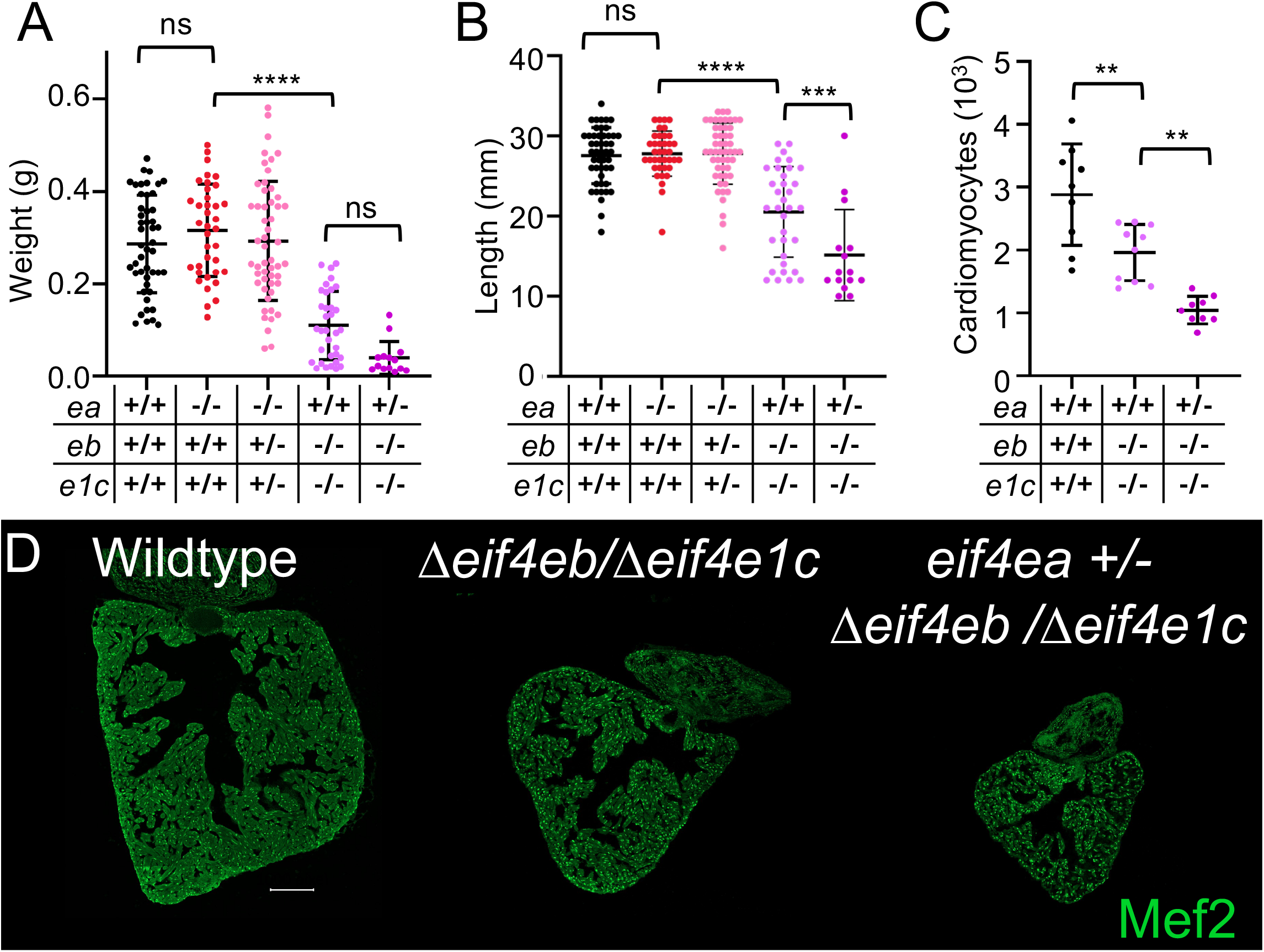
Loss of additional alleles from the genes encoding canonical cap binding factors demonstrates a dose dependence in their compensation abilities. **(A)** Weights of Δ*eif4eb /* Δ*eif4e1c* double mutants compared to of Δ*eif4eb /* Δ*eif4e1c* double / *eif4ea* hets (Mean: wildtype = 0.287g, Δ*eif4ea* = 0.315g, Δ*eif4eb /* Δ*eif4e1c* = 0.110, Δ*eif4ea*-HET /Δ*eif4eb /* Δ*eif4e1c* = 0.039; ANOVA wt - Δ*eif4ea* p-value = 0.712, ANOVA wt-Δ*eif4eb /* Δ*eif4e1c* p value < 0.0001; N = 47, 37, 34, 14). **(B)** Lengths of respective mutants (Mean: wildtype = 27.5mm, Δ*eif4eb /* Δ*eif4e1c* = 20.5mm, Δ*eif4ea*-HET*/*Δ*eif4eb /* Δ*eif4e1c* = 15.1mm, ANOVA p value < 0.001, N = 48, 34, 14). **(C) (**Mean: wildtype = 2882, Δ*eif4eb /* Δ*eif4e1c* = 1962, Δ*eif4ea*-HET*/*Δ*eif4eb /* Δ*eif4e1c* = 1046, ANOVA p value < 0.0001, N = 9, 10, 9). **(D)** Images of representative hearts.

### The canonical cap-binding proteins play differing roles during heart and fin regeneration

Previously, we showed that Δ*eif4e1c* mutants have impaired regeneration of their hearts, suggesting that *eif4e1c* is a pro-regenerative mRNA cap binding protein. The fact that Δ*eif4ea /* Δ*eif4eb* double mutant zebrafish survive in Mendelian ratios gives us the opportunity to determine how hearts regenerate with only *eif4e1c* present. We performed surgeries on Δ*eif4ea /* Δ*eif4eb* double mutant fish and their wild-type siblings and saw that CMs in the double mutant fish proliferate normally (Figure 6AB). Surprisingly, however, the single mutants of either Δ*eif4ea* or Δ*eif4eb* have increased CM proliferation during the peak of heart regeneration (Figure 6AB, Mean: wildtype = 9.6%, Δ*eif4ea* = 12.9%, Δ*eif4eb* = 12.8%, Δ*eif4ea* / Δ*eif4eb* = 8.1%, ANOVA p < 0.0001, N = 48, 37, 40, 38). This suggests that either the canonical cap-binding protein is inhibitory to heart regeneration *or* that an increased ratio of *eif4e1c* is pro-regenerative to the heart. Yet, when both canonical genes are deleted, compensation mechanisms are triggered and the hearts regenerate normally with only *eif4e1c* present.

**Figure 6.**
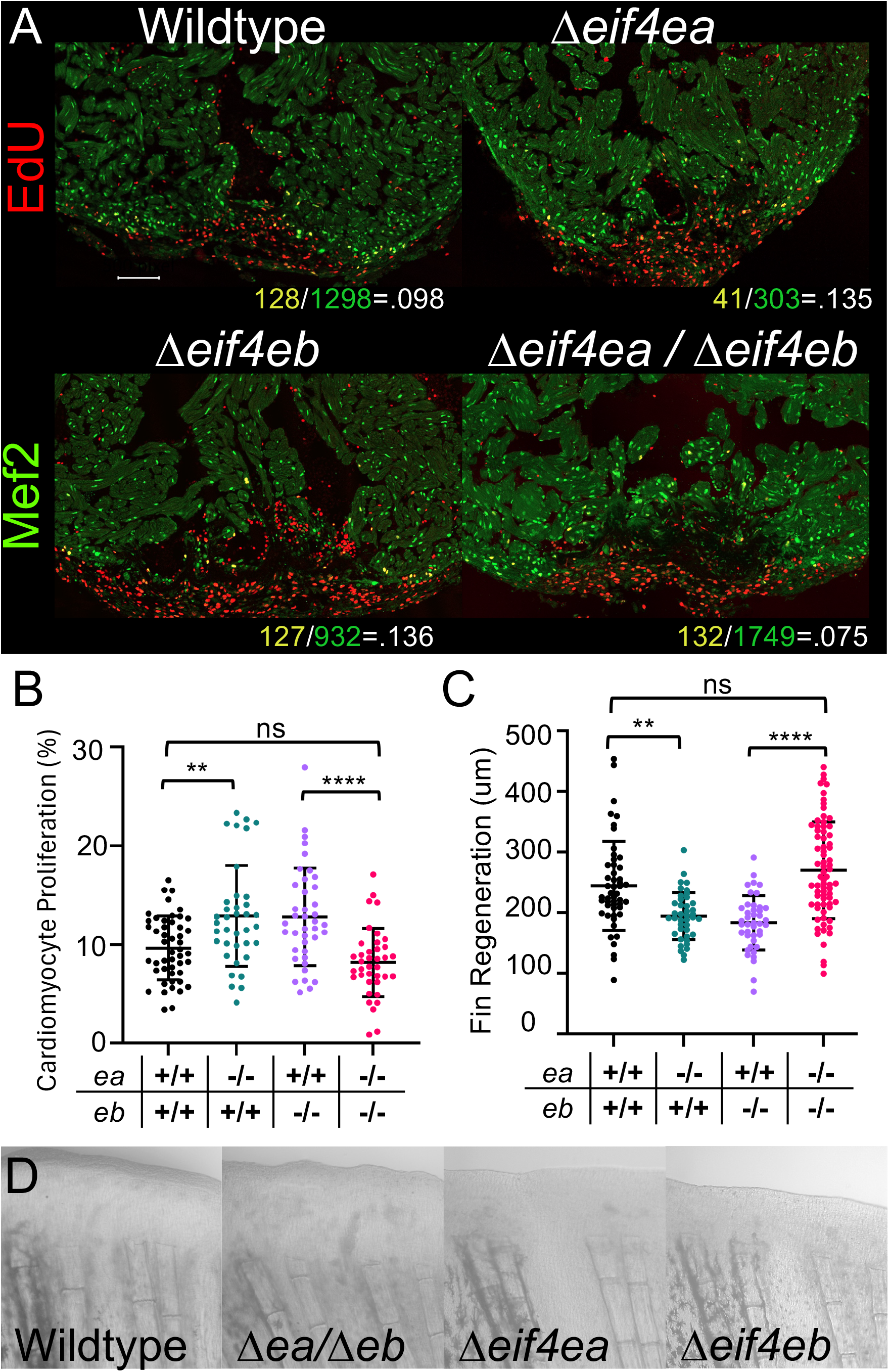
An increased ratio of *eif4e1c* to canonical cap binding alleles improves heart regeneration. **(A)** Hearts, 7 days after amputation of the apex and after 3 days injection of EdU. CM are stained with Δ-Mef2c. **(B)** The single Δ*eif4ea* mutants (green) and the Δ*eif4eb* mutants (purple) have increased levels of CM proliferation (Mean: wildtype = 9.6%, Δ*eif4ea* = 12.9%, Δ*eif4eb* = 12.8%, Δ*eif4ea* / Δ*eif4eb* = 8.1%, ANOVA p < 0.0001, N = 48, 37, 40, 38). However, after a certain threshold, *eif4e1c* fully compensates since Δ*eif4ea /* Δ*eif4ea* double mutants (pink) regenerate like their wildtype (black) siblings. **(C)** Graph of fin regrowth 2 days post amputation in the respective mutants (Mean: wildtype = 244um, Δ*eif4ea /* Δ*eif4eb* = 270um, Δ*eif4ea* = 194um, Δ*eif4eb* = 183um, ANOVA p-value < 0.0001, N= 50, 73, 41, 42). **(D)** Images of representative fins from the graph.

Since mutants in individual canonical genes have an increase in CM proliferation during heart regeneration, we wanted to determine whether canonical cap-binding proteins are also important during regeneration of the fin. We cut caudal fins of Δ*eif4ea /* Δ*eif4eb* double mutants and compared regrowth with wild-type siblings. As in the heart, fin regeneration was not impacted in the double mutants (Figure 6CD, Mean: wildtype = 244um, Δ*eif4ea /* Δ*eif4eb* = 270um, ANOVA p-value = 0.177, N= 50, 73), but phenotypes did emerge for either single Δ*eif4ea* and Δ*eif4eb* mutants (Figure 6CD, Mean: Δ*eif4ea* = 194um, Δ*eif4eb* = 183um, ANOVA p-value < 0.0001, N = 41, 42). However, unlike in the heart, fin regeneration was *decreased* in the individual canonical mutants. Therefore, increasing the ratio of *eif4e1c* versus canonical cap-binding proteins is beneficial to the heart but detrimental to the fin. We conclude that *eif4e1c* specific functions are more crucial to the heart.

## Discussion

Here, we show that the phenotypes we previously reported in Δ*eif4e1c* mutants reflect specialized functions of the *eif4e1c* pathway since they are not recapitulated in mutants for canonical cap-binding factors Δ*eif4ea* and Δ*eif4eb*. However, the phenotypes in Δ*eif4e1c* mutants are incomplete and combination of Δ*eif4e1c* with Δ*eif4ea* or Δ*eif4eb* results in potentiation of phenotypes demonstrating that canonical factors can only partially compensate for *eif4e1c* function. Surprisingly, zebrafish growth and survival are normal in mutants with loss of both paralogs of *eif4ea* and *eif4eb*. This makes the *eif4e1c* pathway unique among the reported mRNA cap-binding homologs because *eif4e1c* can fully compensate for canonical function. While the canonical mutants demonstrate no obvious phenotypes during their normal lifetime, during regeneration, phenotypes emerge. Deletion of either canonical factor *eif4ea* or *eif4eb* results in improved regeneration of the heart but is detrimental to regeneration of the fin. To our knowledge, zebrafish mutants with regeneration phenotypes have only been reported to effect one tissue or if they impact multiple tissues, it is in the same direction. The differential outcomes in the fin and the heart makes the Δ*eif4ea* and Δ*eif4eb* mutants unusual and suggests opposing roles in the two tissues. Taken together, these data suggest a model where Eif4e1c *can* perform both specialized and canonical functions and it is the specialized functions that are beneficial to only regeneration of the heart.

Interestingly, regeneration phenotypes disappear when Δ*eif4ea* and Δ*eif4eb* are combined in a double mutant. Since the individual mutants both show the same regeneration phenotypes, the compound mutants would be expected to have those phenotypes potentiated or at least unchanged. However, canonical mRNA cap binding proteins demonstrate unusual requirements. While some functional protein is required for viability across eukaryotes, their levels can be severely abrogated with little to no obvious phenotypes^16^. For example, in yeast, deletion of eIF4E is lethal, but degradation of the protein by up to 90% results in growth deficits that are subsequently compensated by other pathways in a matter of hours^4,17^. Similarly, in this study, it is only upon total loss of both canonical paralogs that compensation is triggered. Under these conditions, when canonical factors are severely limited, *eif4e1c* canonical abilities predominate over its specialized functions.

One of the major findings of this study is that *eif4e1c* appears to be beneficial to CM proliferation in the heart but the canonical cap binding proteins are more important during fin regeneration. Deletion of a single gene for canonical cap binding proteins improves heart regeneration possibly by raising the ratio of *eif4e1c* relative to canonical factors. While we cannot exclude the alternative hypothesis that the canonical factors are inhibitory, we favor a model where *eif4e1c* is beneficial because previously we showed that the Δ*eif4e1c* mutant has negative consequences for CM proliferation^13^. We showed that the *eif4e1c* pathway is involved with regulating metabolism which is likely more crucial to the heart during regeneration^13,19–21^. Why might the canonical cap-binding proteins be beneficial to fin regeneration? Unlike the heart, fin regeneration proceeds by the formation of a blastema, which is a plane of highly proliferating dedifferentiated cells that appear to have some stem-like capabilities^22^. Expression levels of the canonical cap-binding protein impacts proliferation rates and it is possible that a reduction in cap-binding proteins by half reduces proliferation rates as it has been shown in mice and many cancers^14,18,23,24^. More work remains to be done to untangle mechanistic underpinnings that underlie these alternative regeneration phenotypes.

In this manuscript we demonstrate that genes for both canonical cap-binding protein paralogs partially compensate for Δ*eif4e1c* phenotypes. If either Δ*eif4ea* or Δ*eif4eb* homozygous alleles are combined with Δ*eif4e1c*, then growth is further diminished. However, when it comes to survival, the Δ*eif4ea* and Δ*eif4eb* alleles behave quite differently. The fact that *eif4eb* does not compensate for survival to the same extent as *eif4ea* suggests two ideas. First, that the *eif4ea* and *eif4eb* genes have become specialized and have evolved different properties although both have been found to be expressed in all tissue types and cell-types^13,25^. An ancestor of teleost fish likely had a whole genome duplication almost 400 million years ago and ∼20% of genes have been retained as duplicates with some paralogs reported to have neofunctionalization^26–32^. Second, that survival and growth phenotypes in Δ*eif4e1c* may not be linked to one another and likely result from different *eif4e1c* processes. However, it is possible that the expression levels of the Eif4ea and Eif4eb proteins are dynamic, and this explains the differential compensation. While this idea is not obvious solely by measuring mRNA levels, there may be stage or cell-type specific fluctuations that are not yet appreciated in the published record.

## Materials and methods

### CRISPR mutant production

Pairs of sgRNA fused to tracRNA were assembled with Cas9 and injected into newly fertilized zygotes using the IDT DNA system. For *eif4ea,* sgRNA targeted exon 3 (TGT CAA ACT TTG AGA TGA GA) and exon 7 (CAT GTG ACT GGT ATC CAA TG) that created a 4,262bp deletion. For *eif4eb*, sgRNA targeted exon 2 (GAT CTC AGA CGA GAG CAA TC) and exon 3 (CTC CAA ATT TGA CAC CGT AG) that created a 560bp deletion. Independent lines from different injections were crossed to one another to homozygous mutant alleles and minimize potential for homozygosis of possible off target mutations that would be confounding. We created two lines for *eif4ea*, one with a perfect deletion (L2) and the other that had an insertion of a single G (L1) in place of the deletion. For *eif4eb* we created four lines. Two with *eif4eb* alone and two that were injected into *eif4e1c* carriers to create double mutants since both *eif4eb* and *eif4e1c* are on the same chromosome. For the *eif4eb* solo mutants, one had an insertion of GGATTTTTGGG (L4) and the other deletion was perfect (L5). For the double mutants one allele was perfect (L5) and the second had an insertion of GGG (L8).

### Genotyping and Survival

For genotyping *eif4ea* mutants we used two oligos that bridge the mutation 5’-ACT TGT TCT TGG TGG TGG AAC CGC and 5’-TCT AGA GGA GTT ATC ATC CCC AAC CCC plus a third internal oligo that recognizes the wildtype allele (5’-GGT CAC TTT AGG ATC TGT TCA CCC TGC). For genotyping *eif4eb* mutants we used two oligos that bridge the mutation (5’-TGT TCA AAC TGG CTG CTT TAG TGT GGA and 5’-AGC GAG TAG TCA CAT CCT GAC ATC AG) plus a third internal oligo that recognizes the wildtype allele (5’-GCA AAA CAT GGC AGG CCA ACC). Multiple clutches >50 were combined when considering the Mendelian proportions and data for individual clutches can be found in Table 1.

### Weight and length measurements

Adult zebrafish (3-4 mpf) were anesthetized in phenoxyethanol. Length was measured using a ruler and estimated to the nearest millimeter from the tip of their mouth to the center of the bifurcation of their caudal fin. Fish were then blotted dry with Kimwipes and weighed on an analytical scale. Fish of all genotypes were raised and housed together, and measurements were taken within 2 weeks after genotyping.

### Immunostaining

Hearts were sectioned to 10-12um and placed on slides. In every case possible, we embedded test and control hearts together. For example, either the top or the bottom row was wildtype, and the other row was mutant. Thus, hearts were imaged and analyzed in a blinded fashion and under identical imaging conditions. The Mef2ca antibody was obtained commercially from Boster Biological Technology (DZ01398-1) and used at a 1:400 dilution. Prior to staining, slides were boiled for 10 minutes in citrate acid buffer to unmask epitopes. The block and primary binding buffers were identical (1% BSA, 1% goat serum, 0.025% tween-20). The secondary was anti-rabbit-488 (1:200). Hearts were imaged with a 20X objective using a Zeiss confocal. Images were processed and CMs counted using the MIPAR image processing software^33^.

### Heart regeneration

Adult fish (>5 months) had the apex of their ventricles amputated as previously described^13,34^. Injured fish were injected into the abdominal cavity once every 24 h for 3 days (4-6 dpa) with 10 μl of a 10 mM solution of EdU diluted in PBS. Hearts were removed on day 7, embedded and cryosectioned. Slides were stained with Alexa Fluor 594 Azide using click chemistry^35^ and then immunostained as described above for Mef2c. Heart injury sites were tiled 3 × 2 using the Zeiss confocal and 2-3 different sections were averaged to obtain the final number per heart. Three independent biological replicates were done from three independent clutches (at least 12 fish from each group) that were raised months apart. The final data presented merges the data for the biological replicates.

### Fin regeneration

Caudal fins were amputated at 50% from the tip of the fin to the base of the body. At 2 days post amputation, regrowth was imaged with a Zeiss compound microscope and a 20X objective. The length of regrowth from the amputation plane to the end of the blastema was measured using the ZEN software from Zeiss. Fin regeneration experiments were performed on the same fish that were used for the heart regeneration experiments.

